# Structural basis for sterol sensing by Scap and Insig

**DOI:** 10.1101/2021.06.03.446951

**Authors:** Renhong Yan, Pingping Cao, Wenqi Song, Yaning Li, Tongtong Wang, Hongwu Qian, Chuangye Yan, Nieng Yan

**Affiliations:** Westlake Laboratory of Life Sciences and Biomedicine, Key Laboratory of Structural Biology of Zhejiang Province, School of Life Sciences, Westlake University, 18 Shilongshan Road, Hangzhou 310024, Zhejiang Province, China; Institute of Biology, Westlake Institute for Advanced Study, 18 Shilongshan Road, Hangzhou 310024, Zhejiang Province, China; State Key Laboratory of Membrane Biology, Beijing Advanced Innovation Center for Structural Biology, Tsinghua-Peking Joint Center for Life Sciences, School of Life Sciences, Tsinghua University, Beijing 100084, China; Department of Molecular Biology, Princeton University, Princeton, NJ 08544, USA

**Author notes:** These authors contributed equally to this work (Renhong Yan, Pingping Cao, Wenqi Song, Yaning Li). To whom correspondence should be addressed: N. Yan; C. Yan; R. Yan.

## Abstract

The sterol regulatory element-binding protein (SREBP) pathway senses the cellular cholesterol level through sterol regulated association between Scap and Insig. Despite the recent structural determination of the transmembrane domains of human Scap and Insig-2 bound to 25-hydroxycholesterol (25HC), the structure and regulatory mechanism of the luminal domains of Scap by cholesterol remains elusive. Here, combining cryo-EM analysis and artificial intelligence-facilitated structural prediction, we report the structure of the human Scap/Insig-2 complex in the presence of digitonin instead of 25HC. Despite the lack of sequence similarity, the structure of the luminal domain Loop 1 and a co-folded segment in Loop 7 of Scap resembles that of the luminal/extracellular domain in NPC1 and related proteins. Comparison of the sterol-loaded structures of these proteins provides clues of the regulation of Loop 1/7 interaction by cholesterol. We also show that the structure of Scap(D428A), which suppresses SREBP activation under sterol depletion, is identical to WT when complexed with Insig-2, although the gain of function may also involve a later step in protein trafficking.

## Introduction

Abnormal sterol metabolism and aberrant accumulation of cholesterol contributes to the development of multiple cardiovascular diseases, which have been a major health threat worldwide (*1, 2*). There are two sources for cellular cholesterol, *de novo* synthesis and diet intake. Crosstalk between these two processes and cellular homeostasis of cholesterol is controlled through an end-product feedback mechanism executed by the sterol regulatory element-binding protein (SREBP) pathway (*3–6*).

A critical step in the SREBP pathway is to monitor the cellular sterol level, which is executed by two membrane proteins, the SREBP cleavage-activating protein (Scap) and the insulin-induced gene (Insig-1 or −2). When the cell is replete with cholesterol, Scap binds to the ER-anchored Insig-1 or −2 through its sterol sensing domain (SSD), which is comprised of five transmembrane (TM) segments S2-S6 (*7–9*). As a result, SREBP-2 associates with Scap through their respective cytosolic C-terminal domains and is locked in the ER membrane. When cellular cholesterol drops from certain threshold, the interaction between Scap and Insig cannot be maintained in the absence of the sterol, and the SREBP-2/Scap complex is translocated to the Golgi by the COPII-coated vesicles (*10*). After two steps of proteolytic cleavage, first in the lumen and second within the Golgi membrane, the liberated N-terminal transcription factor domain of SREBP-2 enters the nucleus to activate gene expression for cholesterol synthesis and uptake, restoring the cellular cholesterol level (*11–15*). Therefore, the sterol-regulated interaction between Scap and Insig functions as a switch for the activation or suppression of the SREBP pathway (*6, 16*).

Because of the dysregulation of cholesterol homeostasis in cancer cells and viral infection model, cholesterol metabolism, inhibition of the SREBP pathway connotes a potential strategy for the development of novel therapeutics for cancer treatment and viral defense (*17–21*). Structural elucidation of the SREBP pathway components may thus facilitate drug discovery, as well as provide in addition to affording an advanced mechanistic understanding of the regulation of cholesterol metabolism.

We recently resolved the cryo-EM structure of the human Scap and Insig-2 complex in the presence of 25-hydroxycholesterol (25HC), which is more potent in promoting Scap/Insig complex formation than cholesterol (*22*). In the well-resolved structure of the transmembrane (TM) domains of the complex, 25HC stands at the interface of Scap and Insig-2. Further analysis suggests that the primary mechanism for 25HC to bridge the complex formation may go beyond a simple molecular glue. Insertion of 25HC into a cavity in the luminal leaflet of the complex stabilizes the unwinding of the S4 segment of Scap in the middle, a conformation that is critical for Insig association. In a well-documented mutant, Scap(D428A) (*23*), and a variant rationalized from our structural analysis, Scap(Q432A) (*22*), both modified proteins gained the function of forming complex with Insig in the absence of sterols, supporting the notion that maintenance of an unwound S4 and the nearby S2 segment is sufficient for Insig binding, even without a molecular glue. Nevertheless, the structure of Scap(D428A)/Insig is required to consolidate this analysis.

Analysis of the ternary Scap-Insig-25HC complex reveals a promising interpretation of the 25HC-dependent interaction between Scap and Insig, and an explanation for the greater ability of oxysterols to promote this interaction over cholesterol. Previous studies have suggested a more complicated allosteric regulation mechanism by cholesterol of Scap that involves its luminal domains (*16, 24–26*). It has been shown that cholesterol binding to the luminal domain Loop1 (L1, the segment between S1 and S2) disrupts the interaction between L1 and Loop7 (L7, the segment between S7 and S8) (*24, 25*). The cholesterol-modulated conformational changes lead to the sequestration of the MELADL motif, which marks the C-terminus of S6 on the cytosolic side (*22*), from being recognized by the COPII proteins (*26, 27*). Insufficient resolution of the luminal domains in our previous cryo-EM map prevented model building (*22*). The interaction details of L1/L7 and the molecular basis for their regulation by cholesterol remain to be elucidated.

Here we report the cryo-EM analysis of the Scap/Insig-2 complexes in the absence of 25HC. The cryo-EM structure of WT Scap in complex with Insig-2 reveals that the detergent molecule, digitonin, can replace 25HC to bridge the interaction. Under this condition, a co-folded L1/L7 luminal domain was resolved to adequate resolution and model building was aided by artificial intelligence (AI)-facilitated structure prediction. In addition, our structural and biochemical characterization of the complex of Scap(D428A) Insig-2 variants reveal that Scap(D428A) suppresses SREBP activation even in the absence of Insig binding.

## Results

### Structure of the Scap/Insig-2 complex in digitonin micelles

With the aim of improving the resolution of the luminal domains of Scap, we tested multiple conditions for cryo-EM sample preparation. Similar to our reported protocol (*22*), WD40-deleted human Scap (residues 1-735), WT or D428A, and an Insig-2 variant that contains three point mutations designated Insig-2 3CS (C14S, C90S, C215S) with N terminal Bril fusion, were co-expressed in HEK293F cells. During sample preparation, we found that sterols such as 25HC had to be included until affinity purification to stabilize an intact complex when lauryl maltose neopentyl glycol (LMNG) was used for extraction and purification (*22*). In contrast, when digitonin was used as the primary detergent, the complex survived tandem affinity purifications and size exclusion chromatography (SEC) with a better behavior (Fig. S1A). We set out to solve the complex structure in the digitonin micelles.

The method of cryo-EM data acquisition and image processing was the same as that for 25HC-bound complex purified in LMNG. To improve the resolution, all collected data, including those for WT and the Scap variant containing D428A were initially combined for processing. The combined dataset yielded a map of higher resolution than individually calculated ones, suggesting identical conformations of the two complexes (Fig. S2). We will use Scap/Insig-2D to refer to the two complexes until otherwise indicated.

A 3D EM reconstruction was obtained for the complex at an overall resolution of 4.0 Å in CryoSparc (*28*). A soft mask was applied and refinement of the TM domains led to improved resolutions to 3.3-3.9 Å, highest at the interface (Fig. 1A,B; Figs. S2-4, Table S1). When further refined in Relion 3.0 (*29*), the nominal overall resolution dropped to 4.1 Å, but the soluble region was better resolved than that purified in LMNG and 25HC (designated Scap/Insig-2H), resulting in discernible secondary structural elements (Fig. S2).

**Figure 1.**
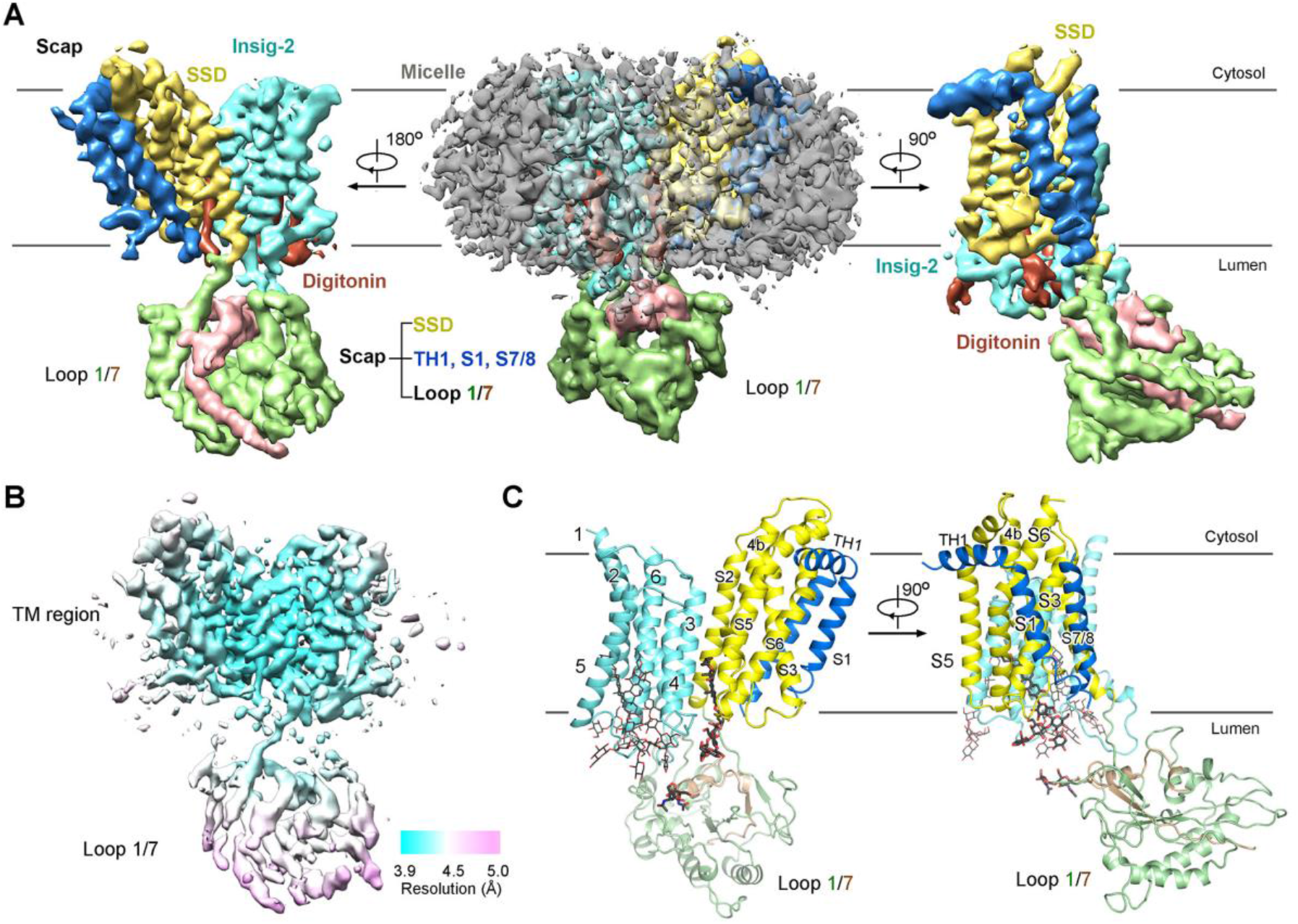
Overall structure of the complex of human Scap and Insig-2 in digitonin micelles. **A**, The 3D EM reconstruction of the complex. The resolution for the 3D EM reconstruction was improved when particles of the Insig-2 complexes with WT and Scap(D428A) were combined, suggesting identical conformation of the two variants when purified in digitonin. Scap/Insig-2D does not distinguish the two complexes if not otherwise indicated. The density is domain colored as annotated. The digitonin micelle is shown as gray, semi-transparent surface to indicate the orientation of the TM domain in the middle panel. The cytosolic side is shown on top for all side views in all figures. The EM maps were generated in Chimera (*42*). **B**, Resolution heat map of the complex. The local densities were calculated using Relion 3.0. **C**, Structure of the human Scap/Insig-2 complex in the presence of digitonin. Two perpendicular side views are shown. Insig-2, the sterol sensing domain (SSD), and the other transmembrane (TM) segments of Scap are colored cyan, yellow, and blue, respectively. The Scap luminal domains Loop 1 and Loop 7 (short as L1/L7) are colored pale green and wheat, respectively. The same color scheme is applied to all figures. The digitonin molecule on the interface of Scap and Insig-2 is shown as black ball-and-sticks and the other three are shown as thin sticks. A glycosylation site on L7 is shown as black sticks. All structure figures were prepared in PyMol (*43*).

In contrast to the 3D EM map for the Scap/Insig-2H complex wherein only one elongated density was observed at the interface between Scap and Insig-2, the present map of the TM domains contains four linear densities, two within and two surrounding Insig-2 (Fig. 1A,C; Fig. S4). Although we cannot exclude the possibility that some of the densities may contain signals contributed by 25HC, which was supplemented at 1 μg/ml during expression and membrane extraction, the four densities all perfectly match digitonin that was used at 1% (w/v) for protein extraction and 0.08% for purification (corresponding to 10 mg/ml and 0.8 mg/ml, respectively).

The structure of 25HC-bound Scap/Insig-2 (PDB code: 6M49) was docked into the EM map for the TM region with manual adjustments, and AI-facilitated structure prediction (*30*) and *de novo* model building were combined for structural assignment of the luminal domains. The details will be described in a later section.

### A potential lateral gate for sterols on the TM interface of Scap and Insig

The overall structures of the TM domains of the complex are similar in the presence of digitonin and 25HC, with a root-mean-square deviation (RMSD) of 1.5 Å over 363 Cα atoms (Fig. 2A). When the two structures are superimposed relative to Insig-2 (Fig. 2B, left) or Scap (Fig. 2C), relatively minor shifts are observed.

**Figure 2.**
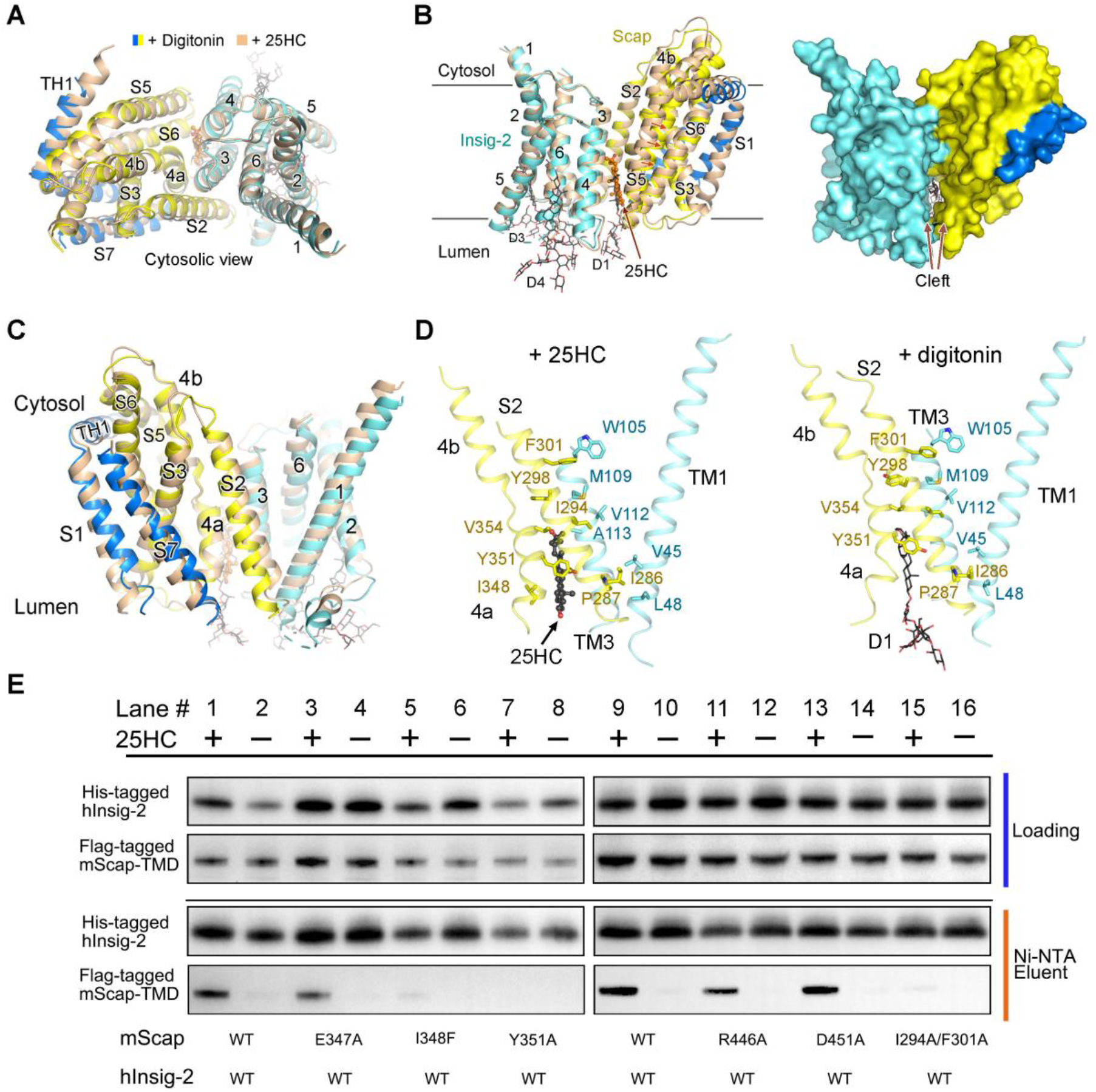
A potential lateral gate for sterols. **A**, Structural comparison of the Scap/Insig-2 complex separately bound to digitonin and 25HC. The TM region of the complex in the presence of digitonin (Scap/Insig-2D) is similar to that bound with 25HC (Scap/Insig-2H, PDB code: 6M49, colored wheat). The two structures can be superimposed with an RMSD of 1.5 Å over 363 Cα atoms. The bound 25HC and digitonin molecules are shown as ball-and-sticks and thin sticks, respectively. **B**, Displacement of Scap from Insig-2 on the S5-TM4 side of the complex results in the formation of a cleft that may represent the lateral gate for sterols. *Left*: Structural shifts of Scap when the two complexes are superimposed relative to Insig-2. Motions of the corresponding segments in Scap from 25HC-bound to the digitonin-purified structure are indicated by orange arrows. *Right*: The interface mediated by Scap-S5 and Insig-TM4 is nearly lost in the presence of digitonin, converting a previously observed fenestration in Scap/Insig-2H (*22*) to a cleft, leaving the sterol binding site open to the lipid membrane. **C**, The direct interface between Scap and Insig-2 is primarily constituted by Scap-S2 and the TM1 and TM3 of Insig-2. The two structures are superimposed relative to Scap, with an RMSD of 1.3 Å over 200 Cα atoms in Scap. **D**, The direct interface between Scap and Insig-2 in Scap/Insig-2D is nearly identical to that in Scap/Insig-2H. Shown here are two identical side views of the complex in the presence of 25HC (*left*) or digitonin (*right*). **E**, Validation of the direct interface between Scap and Insig-2. Ala substitution of Ile348 or Tyr351 in Scap, two residues that are involved in 25HC binding, eliminated 25HC-dependent complex formation. Double Ala mutation of another two Scap residues, Ile294 and Phe301, which mediate the direct interaction between Scap and Insig-2 but away from the 25HC binding site, also completely abolished complex formation. Mutation of Scap residues that are not mapped to the interface, Glu347, Arg446, and Asp451 had little or no influence on 25HC-dependent complex formation. Shown here are representative western blots out of three independent experiments. Please refer to Figure S5 for the original western blot images and Experimental procedures for details.

As shown previously (*22*), the TM interface between Scap and Insig is highly asymmetric. The 25HC-bridged interaction is limited to the luminal leaflet of the membrane on one side involving Scap-S5 and Insig-TM4 (Fig. 2B), while a direct interface that traverses the entire membrane height is mediated by Scap-S2 and TM1 and TM3 of Insig (Fig. 2C,D). Whereas the bound 25HC is insulated from the membrane by this direct interface, a large fenestration forms on the other side wall due to the limited contacts between Scap-S5 and Insig-TM4 (*22*). In the presence of digitonin, which has a bulkier backbone than 25HC, Scap-S5 slightly swings away from Insig-TM4 on the cytosolic side, opening the fenestration to a cleft (Fig. 2B, right). The malleability of the interface between Scap-S5 and Insig-TM4 suggests that this side may represent the lateral gate for sterol entrance and/or exit within the membrane.

Despite the structural shifts of Scap and Insig-2 on one side within the lipid bilayer, the direct interface, which involves multiple hydrophobic residues, remains unchanged in Scap/Insig-2D (Fig. 2C,D). To examine the significance of this interface, we generated a double Ala substitution of Ile294 and Phe301 on Scap-S2 and tested its 25HC-dependent binding with Insig-2. As control experiments, we also engineered several Scap mutants, each containing a single point mutation of residues that may contribute to 25HC binding.

Mutation of either Ile348 or Tyr351 (I348F or Y351A) on S4a, two residues that directly participate in the coordination of the tetracyclic sterane, abolished the association between Scap and Insig-2. Ala substitution of Glu347, which marks the N-terminus of S4a and appears to be in the vicinity of the 3β-OH group of 25HC, only had subtle, if any, effect on the complex formation. Although Ile294 and Phe301 are far away from 25HC, the Scap variant containing double mutations I294A and F301A, lost 25HC-dependent binding to Insig-2, supporting the observed direct interface (Fig. 2E, Fig. S5). In contrast, Ala mutation of residues away from the interface Arg446 or Asp451 had no effect on the complex formation.

### Scap(D428A) exhibits an identical conformation as WT in the complex with Insig

With the reasoning that the potential lateral TM gate between Scap-S5 and Insig-TM4 may undergo more pronounced conformational shifts in the absence of sterols, we sought to solve the structure of the complex between Scap(D428A) and Insig-2. The complex was also co-expressed in HEK293F cells following the same protocol as described above, except for the omission of 25HC throughout protein expression and purification.

Because the structure of Scap(D428A)/Insig-2 is identical to the WT complex in digitonin, we used LMNG for this complex, which survived affinity and SEC purification steps without addition of 25HC (Fig. S1B). This complex, designated Scap(D428A)/Insig-2L, was subject to cryo-EM analysis following the established protocol for WT proteins. In total, 6,347,219 particles were autopicked out of 9,949 micrographs, but only 182,842 particles were selected due to the high heterogeneity. Despite data processing using multiple strategies, the resolution could not go beyond 4.3 Å in contrast to Scap(D428A)/Insig-2D (Fig. S2C).

Although no sterol was added during protein expression and sample preparation for Scap(D428A)/Insig-2L, a weaker density occupies the same position for 25HC in the WT complex (Fig. 3A). We reasoned that this density might belong to endogenous sterols since the cells were not treated for cholesterol depletion during protein expression. We then attempted to use the Sf9 insect cells that have a cholesterol-free background for protein generation. Although this system has been successfully employed for the pull-down assay (*22, 31*), the expression level was too low to support cryo-EM analysis.

**Figure 3.**
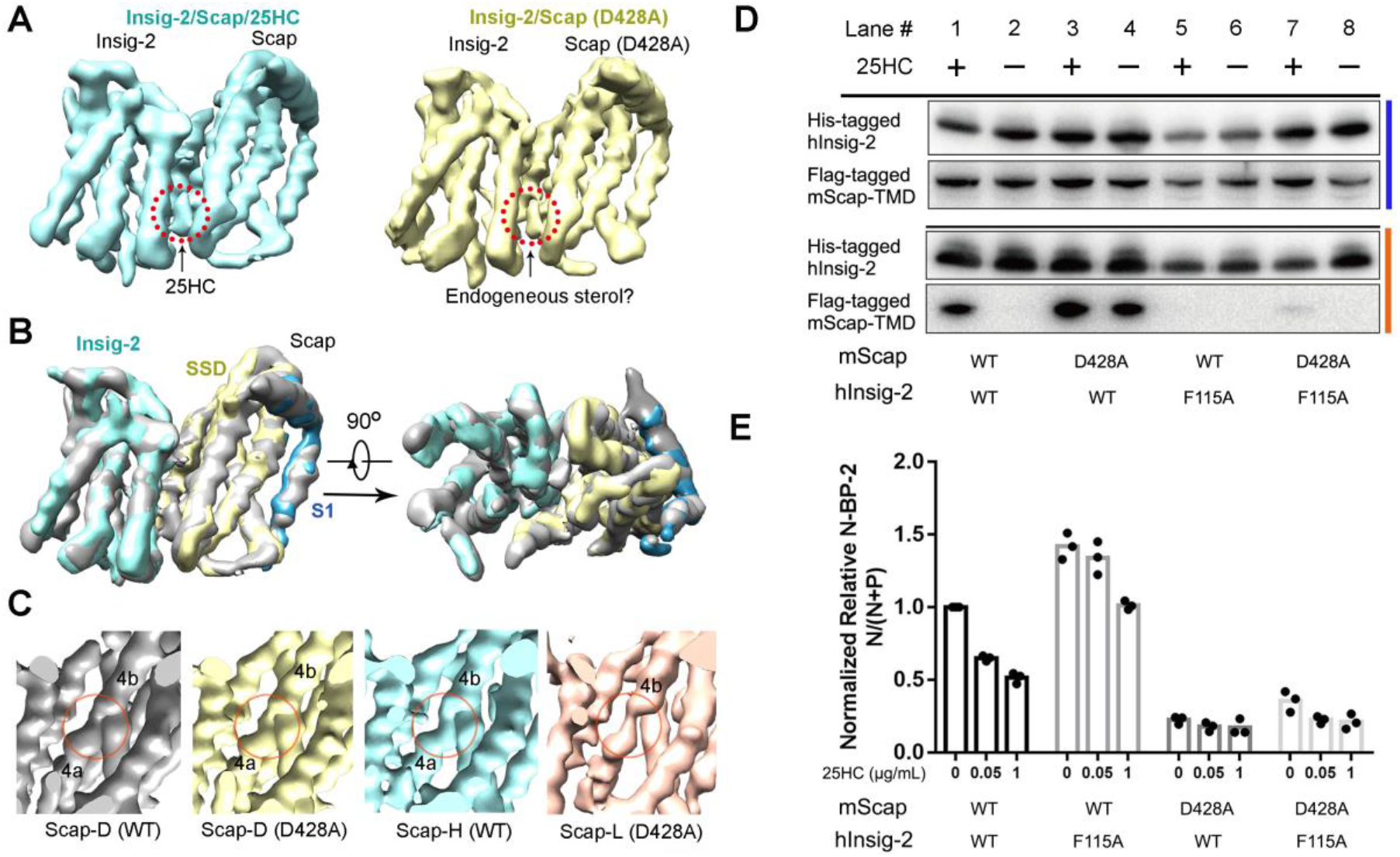
Nearly identical conformations of Scap(D428A) and WT Scap in the complex with Insig-2. **A**, A 3D EM reconstruction of human Scap(D428A)/Insig-2 complex purified exclusively in LMNG was obtained at 4.3 Å. Despite the omission of 25HC throughout expression and purification, a slightly weaker density that may belong to endogenous sterols is observed on the interface between Scap and Insig-2. The two maps were both low-pass filtered to 4.5 Å. **B**, The EM map for Scap(D428A)/Insig-2L (domain colored) is nearly identical to that of Scap(WT)/Insig-2H (grey). Both maps were low pass-filtered to 5 Å. **C**, Scap-S4 is broken in the middle in the Scap(D428A)/Insig-2 complex purified in LMNG or digitonin. Identical views of the EM maps, all low pass-filtered to 4.5 Å, are shown, from left to right, for the WT complex in digitonin, D428A complex in digitonin, WT complex in LMNG and 25HC, and D428A complex in LMNG only. Red circles indicate the unwound region in the middle of Scap-S4 in all four maps. **D**, Scap(D428A) cannot bind to Insig-2(F115A), indicating a mechanism of 25HC-independent loss of function of the latter. Shown here are representative western blots out of three independent pull-down experiments. **E**, Scap(D428A) still suppresses SREBP cleavage when combined with Insig-2(F115A). The loss of binding to Insig-2(F115A) and the inability to activate SREBP cleavage in this context suggest that Scap(D428A) may be trapped in a conformation that is unable to be transported from the ER to the Golgi. The assays were performed in SRD-13A cells transfected with pCMV-3×Myc-mScap and pCMV-His-hInsig-2 variants, as well as the WT pCMV-3×Flag-hSREBP2. Semi-quantification of the band densities was performed using Image J. The mean ratio of the cleaved form over total SREBP2 (N/(N+P)) for WT proteins in the absence of 25HC in three experiments was set as 1, against which the ratio of other groups was normalized. Please refer to Figure S6 for the original western blot images. Details are presented in Experimental procedures.

Although the presence of the sterol-like density prevented us from analyzing the lateral gate, the 3D EM maps of Scap(D428A)/Insig-2 in digitonin and LMNG confirm that Scap(D428A) is well folded and exhibits nearly identical conformation as WT protein, including the broken S4, when complexed with Insig (Fig. 3B,C). This finding is consistent with our previous analysis (*22*).

The sterol-independent association of Scap(D428A) with Insig-2 was thought to account for its gained function of constitutive suppression of the SREBP cleavage. However, we made a serendipitous discovery during our biochemical characterization of Scap and Insig mutants. An Insig-2 variant that contains a single point mutation F115A lost 25HC-dependent binding to WT Scap (*31, 32*) (Fig. 3D, Lanes 1,2,5,6; Fig. S6). Our previous structural analysis suggested that the loss of function may be a result of altered local conformation, independent of 25HC binding (*22*). Supporting this notion, Insig-2(F115A) lost binding to Scap(D428A), with or without 25HC (Fig. 3D, Lanes 3,4,7,8; Fig. S6).

Consistent with the loss of association between Insig-2(F115A) and WT Scap, inhibition of SREBP-2 cleavage by 25HC was substantially reduced in a cell-based SREBP cleavage assay. However, when Scap(D428A) was co-expressed with Insig-2(F115A), cleavage of SREBP-2 was still fully suppressed (Fig. 3E). The contrasting results of the pull-down and SREBP cleavage assays suggest that the constitutive inhibition of the SREBP pathway by Scap(D428A) may not only be attributed to its constitutive association with Insig, but also involve a later step, likely concerning the inter-organellar trafficking.

### Co-folded Loop1 and Loop 7

Next, we examined the luminal domains. Please refer to Experimental procedures for details of model building into this moderate map. Briefly, an AI-predicted model by tFold (https://drug.ai.tencent.com/console/en/tfold) (*30*) for the luminal domain Loop 1 (L1, residues 69-281) of Scap fits the luminal density relatively well (Figs. S7A-C). In particular, the C-terminus of the L1 domain is unambiguously contiguous with S2 in the EM map, serving as a strong structural validation for the assignment of L1 (Fig. S7C). Tracing the backbone suggested that an extra density that intertwines with the assigned L1 could not belong to the invisible residues 47-68. The extra density likely belongs to a segment in Loop 7 (L7). Sequence assignment of L7 (residues 624-661) into this density exhibits a stretched U shape that was supported by large bulky residues and a glycosylation site at Asn641 (Figs. S7D,E). The model was manually adjusted to better fit the density. However, the resolution of the co-folded L1/L7 domain ranges from 3.9 Å to 5 Å. Most side chains of L1/L7 were thereby manually removed to avoid over-interpretation and our analysis was restricted to secondary structural elements only (Fig. 4A).

**Figure 4.**
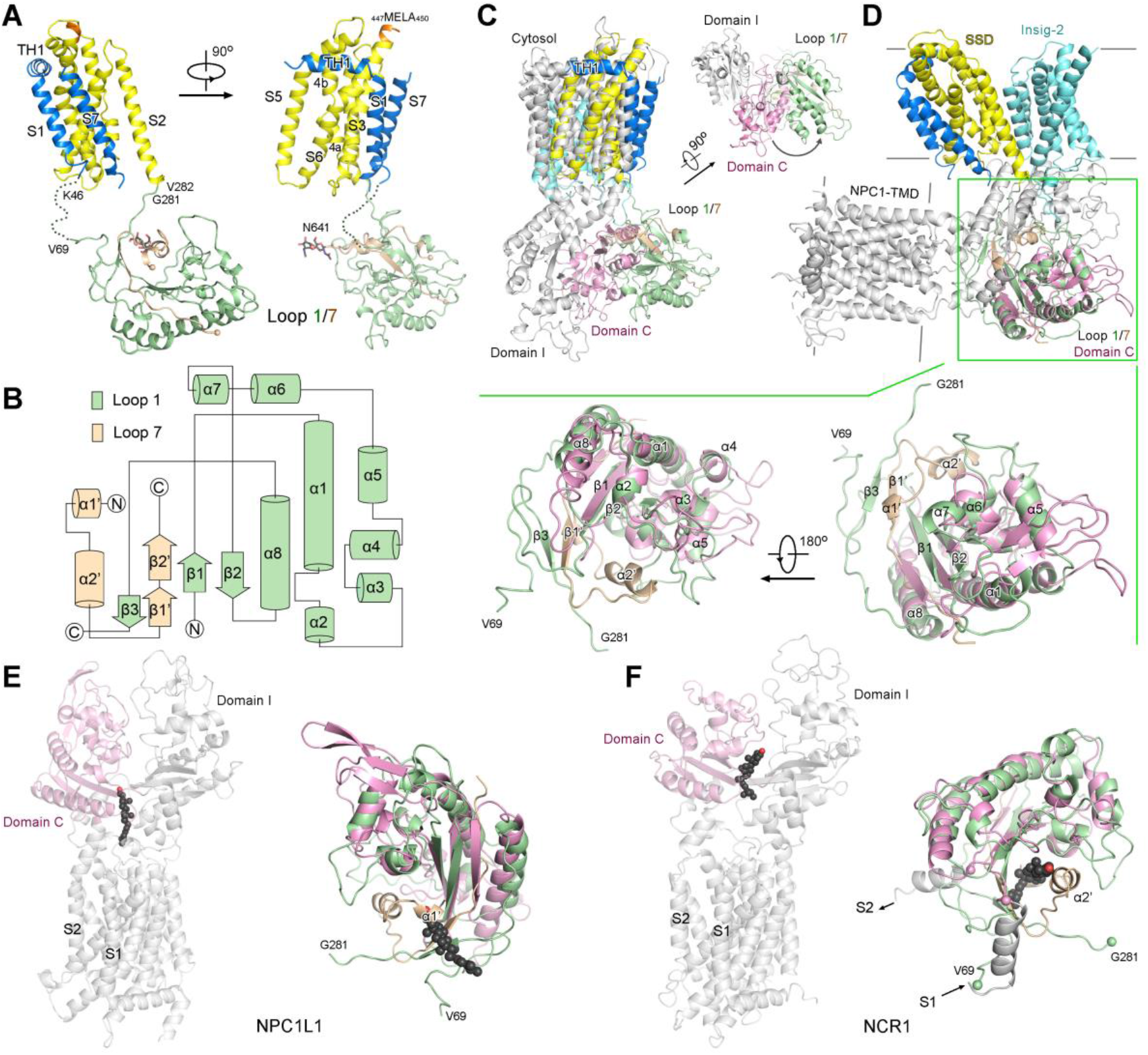
Structural similarity of the Scap-L1/7 domain with the luminal and extracellular domains in other SSD-containing protein. **A**, Overall structure of Scap (residues 1-735) with a moderately resolved luminal domain. The density of the C-terminus of the L1 domain is contiguous with that of the S2 segment, whereas residues 47-68 are invisible on the N-terminus. Only a short segment that co-folds with L1 was resolved in the L7 domain. Please refer to Experimental procedure and Figure S7 for details of AI-facilitated structural modeling. Sequence assignment is facilitated by the glycosylated residue Asn641 and bulky residues. **B**, Topology diagram of L1/L7. **C**, Conformational difference of the luminal domains between Scap and NPC1. When the structure of Scap/Insig-2 complex is superimposed to NPC1 (PDB code: 6W5S) relative to the SSD, the L1/L7 domain of Scap rotates by ~ 80° counterclockwise from domain C of NPC1 in the luminal view. For visual clarity, the N-terminal domain (NTD) and TM0 of NPC1 are omitted. Domain C is colored pink and the rest of the structure is colored silver. The same scheme applies to NPC1L1 and NCR1 shown in panels D & E. **D**, Structural similarity of Scap-L1 and NPC1-domain C despite the lack of obvious sequence homology. Please refer to Figure S8 for sequence alignment of representative SSD-containing proteins. *Upper*: The structure of Scap-L1/L7 is superimposed to that of NPC1-domain C with an RMSD of 2.3 Å over 112 Cα aligned atoms. *Lower*: Enlarged views of the superimposed Scap-L1/L7 and NPC1-domain C. While the L1 domain of Scap comprises an extra β3 strand and additional loops, an elongated segment in L7 overlaps with the last β strand in NPC1-domain C, indicating the co-folding of L1 and L7. **E**, Scap-L7 partially overlaps with a putative cholesterol molecule in the extracellular tunnel of NPC1L1 when the Scap-L1 is superimposed with NPC1L1-domain C. In the structure of NPC1L1 (PDB code: 6V3F), a cholesterol molecule (black spheres) is modeled at the bottom of the extracellular tunnel enclosed by domain C and domain I. **F**, Scap-L7 completely overlaps with the ergosterol bound in the upper half of the luminal tunnel in NCR1 when Scap-L1 is overlaid with NCR1-domain C. The NTD and TM0-deleted structure of NCR1 (PDB code: 6R4L) is shown on the left to indicate the relative position of the ergosterol (black spheres) in the luminal tunnel enclosed by domain C and domain I.

The lower resolution of the luminal domains than the TM domain indicates potential flexibility. Indeed, the resolved L1/L7 domain is featured with large portion of loops, especially the linkers to the TM segments S1 and S2. The core of the luminal domain is comprised of a three-strand β-sheet, β1 and β2 from L1 and β2’ from L7, and two long α helices, α1 and α8 in L1. In addition to the limited sequence for the core, the luminal structure is featured with large number of short helices and long loops (Fig. 4B,C).

We submitted the poly-Ala model of the luminal domains L1/L7 to the Dali server to search for homologous structures (*33*). All the top ranking entries are of SSD-containing proteins, including NPC1 (Niemann Pick disease, type C), NPC1L1 (NPC1-like), NCR1 (NPC1-related), Ptch 1 (patched 1), and Disp (Dispatched) (*34–38*). This result was unexpected because our previous sequence blast failed to reveal sufficient similarities between the luminal/extracellular domains of Scap and the other SSD-containing proteins (Fig. S8). If we had known of this parallel, we would have performed homology-based model building for L1/L7 in our previous structural analysis.

All of the five proteins possess similar architecture including a core structure with 12 TMs, among which S2-S6 form the SSD, and two luminal or extracellular domains. NPC1, NPC1L1, and NCR1 also contain an amino terminal domain (NTD) and an extra TM helix designated by us as TM0. The core structure of these proteins exhibits a 2-fold pseudosymmetry around an axis that is parallel to the membrane norm. The two soluble domains enclose a central tunnel that may serve as the path for cholesterol transport. Densities that correspond to cholesterol or its derivatives have been observed at different positions along the central tunnel (*34–37*).

The structure of L1/L7 most resembles that of domain C in NPC1, NPC1L1, and NCR1, with the RMSD values of 2.3, 3.4, and 2.3 Å over 112, 90, and 113 Cα atoms, respectively, although the overall structures seem different at first sight. When the SSD of Scap is compared to the corresponding region of NPC1, the L1/L7 domain rotates by about 80° counterclockwise from domain C in the luminal view (Fig. 4C). Likewise, when the structures of L1/L7 and domain C are superimposed, the TM domains of the two structures are almost perpendicular to each other (Fig. 4D).

The structures of L1/L7 and domain C are similar in the core region containing all 8 helices in L1 and the β sheet constituted by L1/L7, whereas L1 contains an additional β3 strand, which is connected to the α8 helix through a long linker and forms anti-parallel hairpin with β1’ in L7. This core region of the L1/L7 domain is reminiscent of a half-clenched fist that holds two short helices in L7 (Fig. 4D, bottom insets).

Structural similarity of the soluble domains among the SSD-containing proteins further validates the model building of Scap-L1/L7 and affords mechanistic insight into their regulation by cholesterol. Because of the limitation of the resolution, we will present mechanistic interpretations of the luminal domains in the Discussion session to acknowledge the speculative nature.

## Discussion

### Competition of L1 binding between L7 and cholesterol

Previous studies have shown that cholesterol binding triggers dissociation of L1 and L7, an event followed by the conformational changes of the TM domain for Insig binding and sequestration of the MELADL motif from recognition by COPII proteins (*16, 25–27, 39*). However, the 25HC treatment could not induce the same conformational changes in Scap, suggesting a distinct mechanism for sterol sensing (*40, 41*). Here we show a co-folded structure of the L1/L7 domain of Scap in the presence of digitonin and Insgi-2. Besides, we noticed that the luminal domains of Scap/Insig-2L were of even lower resolution (*22*), indicating higher conformational heterogeneity in the presence of 25HC. Considering that Scap/Insig-2D was isolated and purified in high concentration of the amphiphilic digitonin without treatment of cholesterol, it is possible that digitonin cannot replace cholesterol to disrupt the interaction between L1 and L7. In addition, the complex was deprived of the membrane environment, which may be a prerequisite for the coupling between the luminal domains and the TM segments.

On the other hand, structural comparisons between NPC1L1 and NCR1 (*35, 36*) contain a cholesterol and an ergosterol in the bottom and top of their respective central tunnel, providing some clues of the cholesterol-regulated interaction between L1 and L7 of Scap (Fig. 4E,F). When Scap-L1/L7 is superimposed with NPC1L1-domain C, a short helix in L7 partially overlaps with the bound cholesterol (Fig. 4E). When it is superimposed over NCR1-domain C, the α2’ helix in L7 and the β1’-β3 hairpin completely occupy the position of the bound ergosterol (Fig. 4F). This comparison suggests that cholesterol may compete with L7 to be held by the “palm” of L1.

### A potential luminal interface between Scap and Insig

The better resolved luminal region also allows for modeling of the connecting loop between TM1 and TM2 of Insig-2, designated the L1-2 loop, which contains several bulky side chains. Interestingly, the L1-2 loop appears to project into a cavity enclosed by segments from both L1 and L7 (Fig. 5A, Fig. S7F). Because the entrance to the central pocket of Insig-2 is filled by the bulky sugar moieties in the digitonin molecules, we previously thought that the L1-2 conformation represented an artificial one. However, when we lowpass-filtered the map of Scap/Insig-2H and Scap(D428A)/Insig-2L, similar conformation of L1-2 was observed to contact the luminal domain (Fig. 5B).

**Figure 5.**
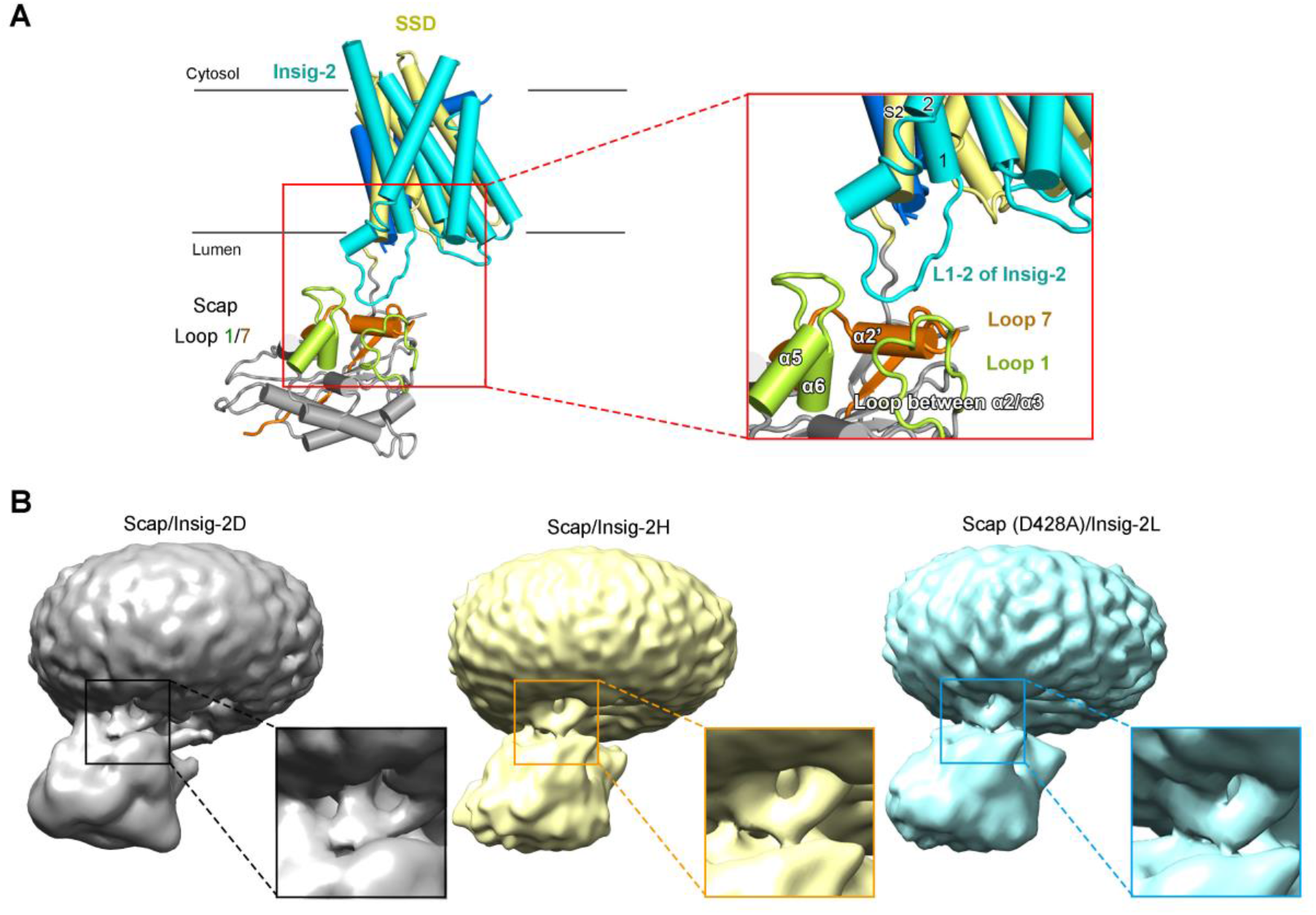
A potential luminal interface between Scap and Insig. **A**, The L1-2 loop of Insig-2 appears to interact with both L1 and L7 of Scap. Please refer to Figure S7F for the density corresponding to the L1-2 loop in Insig-2. **B**, The contact between the L1-2 loop of Insig-2 and the luminal domain of Scap is observed in all three 3D EM reconstructions. Shown here are cryo-EM maps of the indicated complex lowpass filtered to 8 Å.

### Accommodation of digitonin molecules in Insig-2

An outstanding enigma of our previous structure of Scap/Insig-2H is the role of the central cavity of Insig, which is empty in the EM reconstruction (*22*). In the present structure of Scap/Insig-2D, the pocket is filled with digitonin. For illustration simplicity, we designate the digitonin molecule on the protein interface as D1, the two in the central pocket of Insig-2 as D2 and D3, and the last one on the surface constituted by TM3/5/6 as D4 (Fig. 6A).

**Figure 6.**
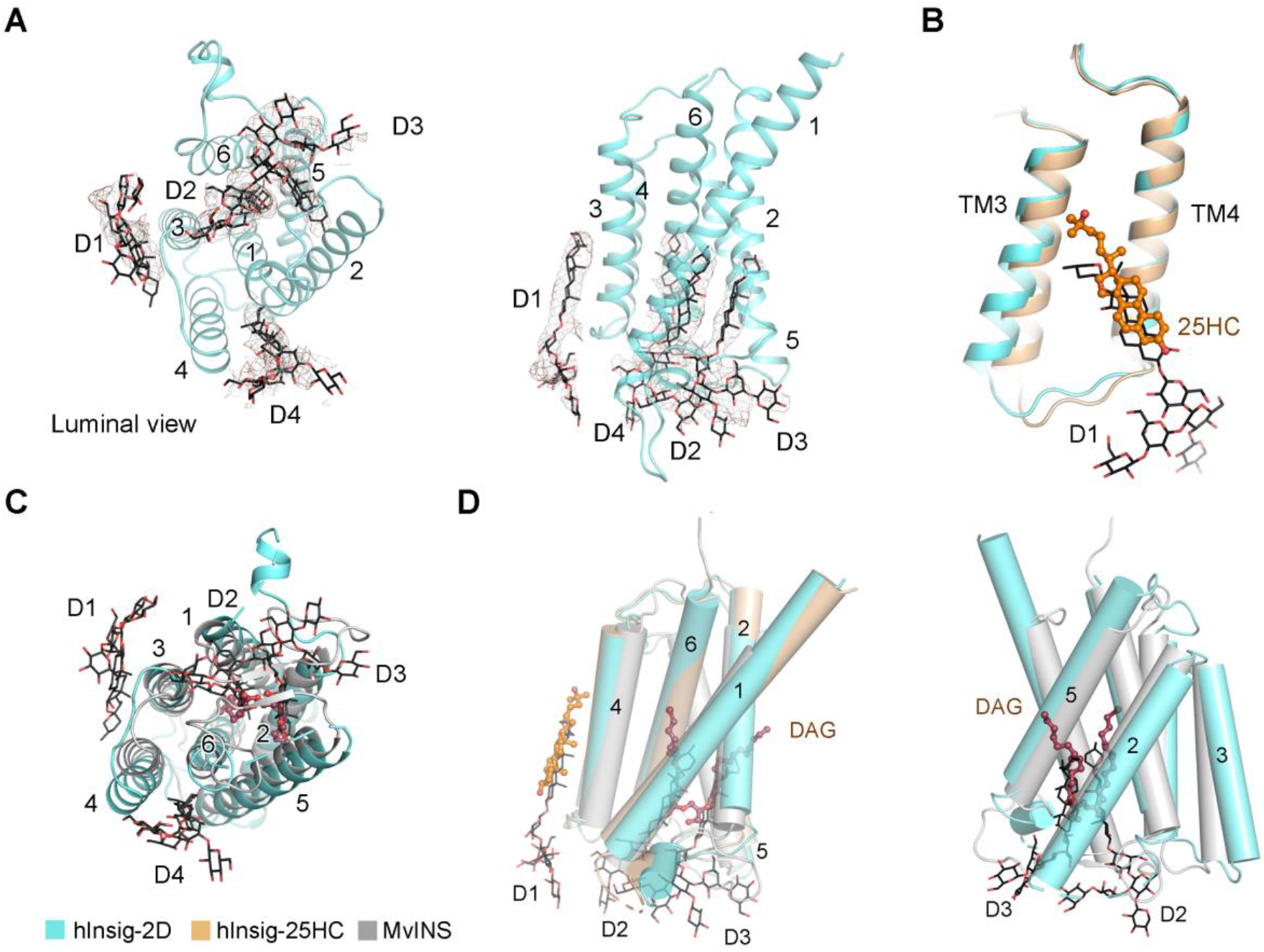
Multiple digitonin molecules bind to the central pocket and lateral surface of Insig-2. **A**, Four stretches of elongated densities bound to Insig-2. The densities that likely belong to digitonin molecules are shown as brown meshes and contoured at 5 σ. The digitonin molecules that bind to the interface between Insig-2 and Scap, in the central pocket, and on the other surfaces of Insig-2 are designated D1, D2, D3, and D4, respectively. **B**, The D1 digitonin molecule overlaps with 25HC. Structures of Insig-2 in the two complexes can be superimposed with an RMSD of 1.0 Å over 155 Cα atoms. The structure in the presence of 25HC (hInsig-25HC) is colored light brown. **C**, Structural comparison between human Insig-2 and MvINS, a mycobacterial homologue of Insigs (*31*). The two structures are overlaid with an RMSD of 3.3 Å over 141 Cα atoms. MvINS: Insig from *Mycobacterium vanbaalenii* (PDB code: 4XU4). **D**, Two digitonin molecules in the pocket of human Insig-2 overlap with the two tails of DAG. *Left*: Superimposition of the two Insig-2 structures and MvINS. D4 digitonin is omitted for clarity. *Right*: When the structures of Insig-2D and MvINS are overlaid, the two digitonin molecules, D2 and D3, in hInsig-2D overlap with the two aliphatic tails of DAG (brown ball-and-sticks) in MvINS.

While D1 well overlaps with 25HC when the two structures are superimposed relative to Insig-2 (Fig. 6B), D2 and D3 are not randomly positioned. In MvINS, a central pocket is connected to the cleft between TM2 and TM5 to cast a reverse V-shaped contour that accommodates a density reminiscent of diacylglycerol (DAG) (*31*). In Insig-2, the central pocket also has a similar lateral cleft. Our previous analyses confirmed that blockage of the central pocket did not affect 25HC-dependent Scap association or SREBP cleavage suppression (*22*). Intriguingly, D2 and D3 in Insig-2D overlap with the two acyl chains of DAG when the two structures are superimposed (Fig. 6C, D). It remains to be investigated whether Insig can accommodate other ligands, such as DAG or phospholipids, for moonlighting functionality is yet to be characterized.

In summary, the structures of the human Scap/Insig-2 complex reported here and previously (*22*) have addressed several key mechanistic questions on sterol sensing in the SREBP pathway. The structures also represent a starting point for emerging questions, such as the physiological relevance of the potential interface between the L1-2 loop of Insig and L1/L7 of Scap and the function of the Insig central pocket. Furthermore, the unexpected structural similarity of the Scap luminal domains with that in other SSD-containing proteins provides some clues to the regulation of the L1/L7 interaction by cholesterol. Addressing this question entails high resolution structures of Scap in the presence of cholesterol. In addition, structure of Scap(D428A) alone is required to elucidate the mechanism of its constitutive SREBP-suppressing activity. Last but not least, inhibition of the SREBP pathway through retention the Scap/SREBP complex from trafficking to the Golgi by enforcing the interaction between Scap and Insig or interfering with the recognition of Scap by the COPII proteins may afford potential therapeutic opportunities for the treatment of cancer and defense against viral infection. Our structural revelation of the complex in the absence of 25HC may facilitate drug discovery towards this goal.

## Acknowledgements

We thank the Cryo-EM Facility and the supercomputer center of Westlake University and the Tsinghua University Branch of China National Center for Protein Sciences (Beijing) for providing cryo-EM and computation support. We thank Xiaomin Li (Tsinghua University) for technical support during EM image acquisition. We thank Amari Tankard in Princeton University for critical reading of the manuscript. This work was funded by the National Key R&D Program of China (2020YFA0509301 to C.Y.) from Ministry of Science and Technology of China, Beijing Nova Program (Z201100006820039 to C.Y.), the China Postdoctoral Science Foundation (2020M681937 to R.Y), and the National Postdoctoral Program for Innovative Talents of China (BX20200304 to R.Y). N.Y. acknowledges the Ministry of Science and Technology of China and the National Natural Science Foundation of China for multiple funds that have supported this research in 2008-2017. N.Y. has been supported by the Shirley M. Tilghman endowed professorship from Princeton University since 2017.

## Author contributions

N.Y. conceived and supervised the project. N.Y., R.Y., P.C., W.S., H.Q., and C.Y. designed the experiments. R.Y., Y.L., P.C. and W.S. performed molecular cloning, protein purification, and cryo-EM data acquisition. P.C., W.S., R.Y., Y.L., and T.W. performed all functional assays. C.Y. performed data processing and structural building and refinement. All authors contributed to data analysis. N.Y. and C.Y. wrote the manuscript.

