## Supplemental Figures S1-S8 and Table S1 for "Structural basis for sterol sensing by Scap and Insig"

**EXPERIMENTAL PROCEDURES**

**Protein expression and purification**

The protocol for protein co-expression is identical to our recently reported one (*1*). The cDNAs of full-length human Insig-2 (Accession: NM_001321329) wild type (WT) or 3CS mutant (C14S, C90S, C215S) and WD40-deleted human Scap (Accession: NM_012235, residues 1-735) were subcloned separately into the pCAG vector. Scap was tagged with an N-terminal FLAG, and the N-terminus of Insig-2 was tandemly fused with 10×His and Bril. The single point mutations were introduced using a standard two-step PCR.

HEK 293F cells (Invitrogen) were cultured in SMM 293T-I or SMM 293T-II medium (Sino Biological Inc.) at 37 °C under 5% CO_2_ in a Multitron-Pro shaker (Infors, 130 rpm). When the cell density reached 2.0×10^6^ cells per ml, the cells were transiently co-transfected with the two plasmids and polyethylenimines (PEIs) (Polysciences). For transfection, two plasmids, ~ 0.75 mg of each were premixed with 3 mg PEIs in 50 mL fresh medium, which, after incubation for 15 mins, was added into one liter of cell culture, and incubated with the cells for 15 mins. 24 hours after transfection, 1 μg/ml 25-hydroxylcholesterol (25HC, Sigma) was added into the cell culture co-transfected with wild-type (WT) Scap and Insig-2(3CS). The transfected cells were cultured for 48 hours before being collected and resuspended in the buffer containing 25 mM Tris (pH 8.0), 150 mM NaCl and a protease inhibitor cocktail containing aprotinin at 1.3 μg/ml, pepstatin at 0.7 μg/ml, and leupeptin at 5 μg/ml.

To purify the complex between WT Scap (residues 1-735) and Bril fused Insig-2 (3CS) in the presence of digitonin (Scap/Insig-2D), the resuspended lysate was solubilized at 4 °C for 2 hours with 1% (w/v) digitonin (Sigma) and 1 μg/ml 25HC. After centrifugation at 18,700×g for 1 hour, the supernatant was collected and loaded onto anti-FLAG M2 affinity resin (Sigma). The resin was rinsed with the buffer (W1 buffer) containing 25 mM Tris (pH 8.0), 150 mM NaCl, 0.08% digitonin. The protein was eluted with W1 buffer supplemented with 0.2 mg/ml FLAG peptide. The eluent was then applied to nickel resin (Ni-NTA, Qiagen). After rinsing with W1 buffer supplemented with 10 mM imidazole, the protein complex was eluted from the nickel resin with W1 buffer supplemented with 300 mM imidazole. The eluent was then concentrated and subjected to size-exclusion chromatography (SEC, Superose 6 Increase 10/300 GL, GE Healthcare) in the buffer containing 25 mM Tris (pH 8.0), 150 mM NaCl, and 0.08% digitonin. The peak fractions were pooled and concentrated for EM analysis.

For the Scap (residues 1-735, D428A) and Bril fused Insig-2 (3CS) complex in digitonin, designated as Scap(D428A)/Insig2D, the protocol was identical except that 25HC was omitted for the whole process. Purification of the complex between Scap (residues 1-735, D428A) and Bril fused Insig-2 (WT) in lauryl maltose neopentyl glycol (LMNG), designated as Scap(D428A)/Insig2L, was the same as described previously (*1*) except for the omission of 25HC during protein expression and purification.

***In vitro* pull-down of mScap by Insig-2**

The pull-down assay was performed following our published protocol (*2*). Human Insig-2, WT or indicated variants, with an N-terminal His_10_-tag and WD40-deleted mouse Scap (residues 1-741), WT or indicated variants, with an N-terminal FLAG tag were co-expressed using baculovirus-infected Sf-9 insect cells, which were cultured in the SF-900 II medium (Gibco) supplemented with 100 units/ml penicillin and 100 μg/ml streptomycin at 27 °C. Where indicated, 25HC, dissolved in ethanol, was added to the culture at indicated concentrations 24 hours after infection. Otherwise, 25HC was omitted throughout the experiments. Cells were collected and resuspended in the TBS buffer (25 mM Tris (pH 8.0) and 150 mM NaCl) plus protease inhibitor cocktail, 1% LMNG, and 0.1% (w/v) cholesteryl hemisuccinate Tris salt (CHS) 40 hours after infection. 25HC was added at indicated concentrated where it was required. After incubation at 4 °C for 1 hour, the mixture was applied to centrifugation at 18,000×g at 4 °C for 30 mins and the supernatant was applied to Ni-NTA. The resin was washed with 1 ml TBS buffer plus 30 mM imidazole, 0.01% LMNG, and 25HC at indicated concentration for seven times. The proteins were eluted using wash buffer plus 300 mM imidazole, applied to 12% NuPAGE (Invitrogen), and analyzed by western blot.

For western blotting, the proteins were transferred to Immobilon-P transfer membranes (Millipore) after SDS-PAGE. The membranes were blocked by 5% (w/v) Bovine Serum Albumin (Amresco) at room temperature for 1 hour and incubated with the primary antibodies – anti-His tag mouse monoclonal antibody (CWBIO) for Insig-2 or anti-Flag mouse monoclonal antibody (CWBIO) for Scap – at 4°C overnight with a 1:2000 dilution. Then the membranes were washed with the TBST that contained 25 mM Tris (pH 8.0), 150 mM NaCl, and 0.05% (w/v) Tween-20 for four times. The bound antibodies were visualized by chemiluminescence (SuperSignal West Dura Extended Duration Substrate; Thermo) using a 1:3000 dilution of goat anti-mouse IgG antibody, Horseradish Peroxidase (HRP) conjugates, as the secondary antibody (CWBIO). Membranes were exposed by ChemiDOC XRS+ System (Bio-Rad).

**SREBP-2 cleavage assay**

On day 0, SRD-13A cells, a Scap-deficient cell line derived from CHO-7 cells (*3*), were distributed in 6-well plates at a density of 2.5×10^5^ cells per well in medium A that contained DMEM/F12 (1:1, Gibco) supplemented with 5% newborn calf serum (Gibco), 5 µg/ml cholesterol (Sigma), 1 mM sodium mevalonolactone (Sigma), and 20 µM sodium oleate (Sigma). After incubation at 37°C for 1-2 h, cells in each well were transfected with plasmids as following: 0.8 µg pCMV-3×Myc-mScap (mouse) variants, 0.9 µg pCMV-His-hInsig-2 (human) variants, and 0.09 ug pCMV-3×Flag-Hsrebp-2 (human) by ViaFect^TM^ transfection reagent (Promega). The medium was removed 16-20 hours later, and cells were washed twice with PBS that contained 135 mM NaCl, 4.7 mM KCl, 10 mM Na_2_HPO_4_, and 2 mM NaH_2_PO_4_ (pH 7.3±0.1) before changing for medium B (medium A supplemented with 5% lipoprotein-deficient serum, 50 µM mevastatin, and 50 µM sodium mevalonolactone) plus 1% hydroxypropyl-β-cyclodextrin (Sigma). After incubation at 37°C for 1 h, cells were washed twice with PBS and switched to medium B in the absence or presence of 25-HC. Protease inhibitors, 1.5 µM FMK-Z-VAD (MedChemExpress) and 20 µg/mL N-acetyl-leucinal-leucinal-norleucinal (ALLN) (MedChemExpress), were added to prevent degradation of the N-terminal fragment of SREBP-2. After incubation for another 4-5 hours at 37℃, cells were washed twice with cold PBS and subsequently suspended in 100 µL lysis buffer (PBS supplemented with 0.5% SDS, protease inhibitors, and benzonase). The cell lysates were left on a shaker at 4°C for 15 min. The total protein concentration of each cell lysate was determined by UV280 using Nanodrop (Thermo Fisher). The cell lysates were mixed with 5 × sample buffer (250 mM Tris-HCl (pH 6.8), 10% SDS, 50% glycerol, 0.2% bromphenol blue, and 500 mM dithiothreitol). Then an aliquot (10~20 µg total protein) of each sample was loaded to NuPAGE^TM^ 4-12% Bis-Tris gradient Gel (Invitrogen). After electrophoresis, the proteins were transferred to Hybond-C nitrocellulose membranes (Bio-Rad). Strips were blocked with 4% (w/v) milk in TBST, and incubated with primary antibodies for 1 h at room temperature. The following antibodies were used: anti His-tag mouse monoclonal antibody (CWBIO), anti-Flag-tag mouse monoclonal antibody (MBL International), and anti c-Myc tag mouse monoclonal antibody (9E10, Invitrogen). After incubation with the primary antibodies, the washed membranes were incubated with HRP conjugated goat anti-mouse secondary antibodies (CWBIO) at room temperature for 1 h. Membranes were exposed by ChemiDOC XRS+ System (Bio-Rad). The band intensities of precursor (P) and nuclear (N) form of SREBP-2 were measured by densitometry using the software ImageJ. Resulting data were analyzed by GraphPad Prism 7.00. The mean ratio of the cleaved form to total SREBP-2 (N/(N+P)) for WT proteins in the absence of 25HC in three experiments was set as 1, against which the ratio of other groups was normalized.

**Cryo-EM sample preparation and data acquisition**

Each purified complex was concentrated to approximately 10 mg/ml. Aliquots (3.5 μl each) of the protein complex were placed on glow-discharged holey carbon grids (Quantifoil Au R1.2/1.3, 300 mesh), which were subsequently blotted for 3.0 s or 3.5 s and flash-frozen in liquid ethane cooled by liquid nitrogen with Vitrobot (Mark IV, Thermo Fisher Scientific). The prepared grids were transferred to a Titan Krios operating at 300 kV equipped with Gatan K2 Summit detector, or Gatan K3 Summit detector for samples purified in digitonin or LMNG, and a GIF Quantum energy filter. Each stack was aligned and summed using the whole-image motion correction program MotionCorr (*4*) or MotionCor2 (*5*).

**Data processing**

For structural analysis of the complex in digitonin, the data for Scap (WT)/Insig-2D and Scap (D428A)/Insig-2D were initially combined in the hope to increase resolution. A total of 7,490,749 particles were automatically picked by Gautomatch (developed by Kai Zhang, [http:/www.mrc-lmb.cam.ac.uk/kzhang/Gautomatch/](http://www.mrc-lmb.cam.ac.uk/kzhang/Gautomatch/)) from 8563 micrographs of Scap (WT)/Insig-2D complex and 7118 micrographs of Scap (D428A)/Insig-2D. Two rounds of 2D classification and three rounds of guided multi-reference classification were then performed using CRYOSPARC (*6*), resulting in 472,221 particles with pixel size 2.18 Å (Fig. S2A,B). Details of this modified procedure of guided 3D classification was previously described in the manuscript reporting the cryo-EM structure of the human spliceosomal C^*^ complex (*7*). The cryo-EM map of Scap/Insig-2H complex (EMDB: EMD-30074) and two additional bad references were low-pass filtered to 20 Å as the initial references. These selected particles were then re-centered and re-extract with pixel size 1.09 Å (Fig. S2C). After three additional rounds of guided multi-reference classification, remaining 252,929 particles yielded a reconstruction at an average resolution of 4.0 Å after Non-Uniform refinement (Figs. S2C, 3A), and that of the TM region was improved to 3.7 Å after focused refinement. The final 252,929 particles were then split into two datasets based on the initial input: Scap (WT)/Insig-2D and Scap (D428A)/Insig-2D, which were separately refined to 4.2 and 4.4 Å, respectively (Fig. S3B), with the initial reference low pass filtered to 20 Å. There is no detectable conformational change between the two reconstructions. To further improve the quality of cryo-EM density in luminal region, we optimized the calculation with RELION 3.0 (*8, 9*), which yielded an overall 4.1 Å resolution map with improved L1/L7 domain (Figs. S2C, 3D; Table S1). The 4.1 Å reconstruction was used for model building and refinement.

For the Scap (D428A)/Insig-2L dataset, a similar procedure was applied and 182,842 good particles yielded a reconstruction at 4.3 Å for the TM region (Figs. S2B, 2C, S3A; Table S1).

The angular distributions of the particles used for the final reconstruction of the Scap/Insig complex are reasonable (Fig. S3C), and the refinement of the atomic coordinates did not suffer from severe over-fitting (Fig. S3E). The resulting density maps display clear features for the secondary structural elements and side chains of the Scap/Insig complexes in the core region.

Reported resolutions were calculated on the basis of the FSC 0.143 criterion, and the FSC curves were corrected with high-resolution noise substitution methods (*10*). Local resolution variations were estimated using RELION 3.0 (*8, 9*).

**Model building and refinement**

Due to the resolution limitation, the atomic coordinates for the Scap/Insig-2D complex were generated by combining homology modelling, structure prediction by artificial intelligence (AI), and *de novo* model building (*11*).

The structure model for TM region of Scap and Insig-2 (PDB code: 6M49) was docked into the density map and manually adjusted in COOT (*12*). Four digitonin molecules were then identified in the TM region after amino acid residue assignment.

A predicted model for L1 (residues 69-281) of Scap was generated by the protein structure prediction server iDrug (https://drug.ai.tencent.com/console/en/tfold) (Fig. S7A). After removing the unmatched N-terminal and C-terminal region of the predicted structure based on the map, residues 90-260 region of L1 was refined into the density (Fig. S7B). Fortunately, the C-terminus (resides 261-280) of the L1 domain is unambiguously contiguous with the well-resolved S2 in the EM map. After the backbone tracing of most sequence of L1, a strip of extra density remains in the middle of the luminal region, likely belonging to L7 or the N-terminal of L1 (Fig. S7D). If the density had belonged to the N-terminal of L1, the segment (residues 48-89) would have to form an internal knot, and the length was insufficient to cover the entire density. Therefore, this strip most likely belongs to L7. Sequence analysis of L7 (residues 541-709) identified residues 624-661 to best fit the density because of the pattern for bulky residues, such as His632, Trp633, Phe637, Tyr 639, and Tyr640, and more importantly, a glycosylation site at Asn641 (Fig. S7E). The resolution of the co-folded L1/L7 domain ranges from 3.9-5 Å. To avoid over-interpretation of the structural model, most side chains of L1/L7 were then removed in the final model.

The Scap/Insig-2D model was refined against the 4.1-Å map using PHENIX (*13*) in real space with secondary structure and geometry restraints. Overfitting of model was monitored by refining the model in one of the two independent maps from the gold-standard refinement approach, and testing the refined model against the other map (*14*) (Fig. S3E). The structures of the Scap/Insig complex were validated through examination of the Molprobity scores and statistics of the Ramachandran plots (Table S1). Molprobity scores were calculated as described (*15*).

**Supplemental Figures and Legends**

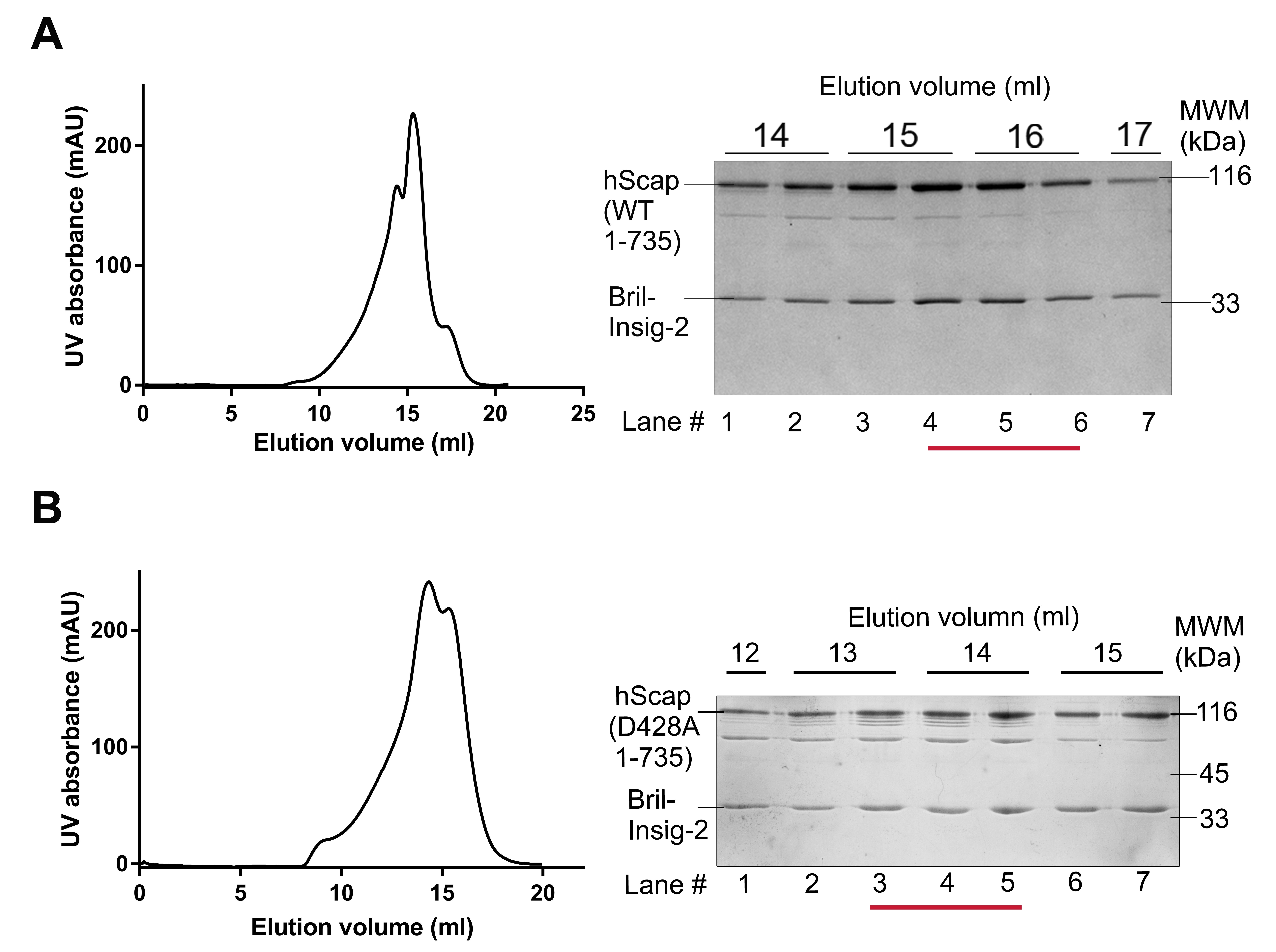

**Fig. S1 |** **SEC purification of the Scap/Insig-2 complex.**

(**A**) SEC purification of the human Scap (WT)/Insig-2 (3CS) complex in 0.08% digitonin. Fractions in Lanes 4-6 were pooled for cryo-sample preparation. Details are presented in Methods. (**B**) SEC purification of the human Scap (D428A)/Insig-2 (WT) complex in LMNG. Fractions in Lanes 3-5 were pooled for cryo-sample preparation. Details are presented in Methods.

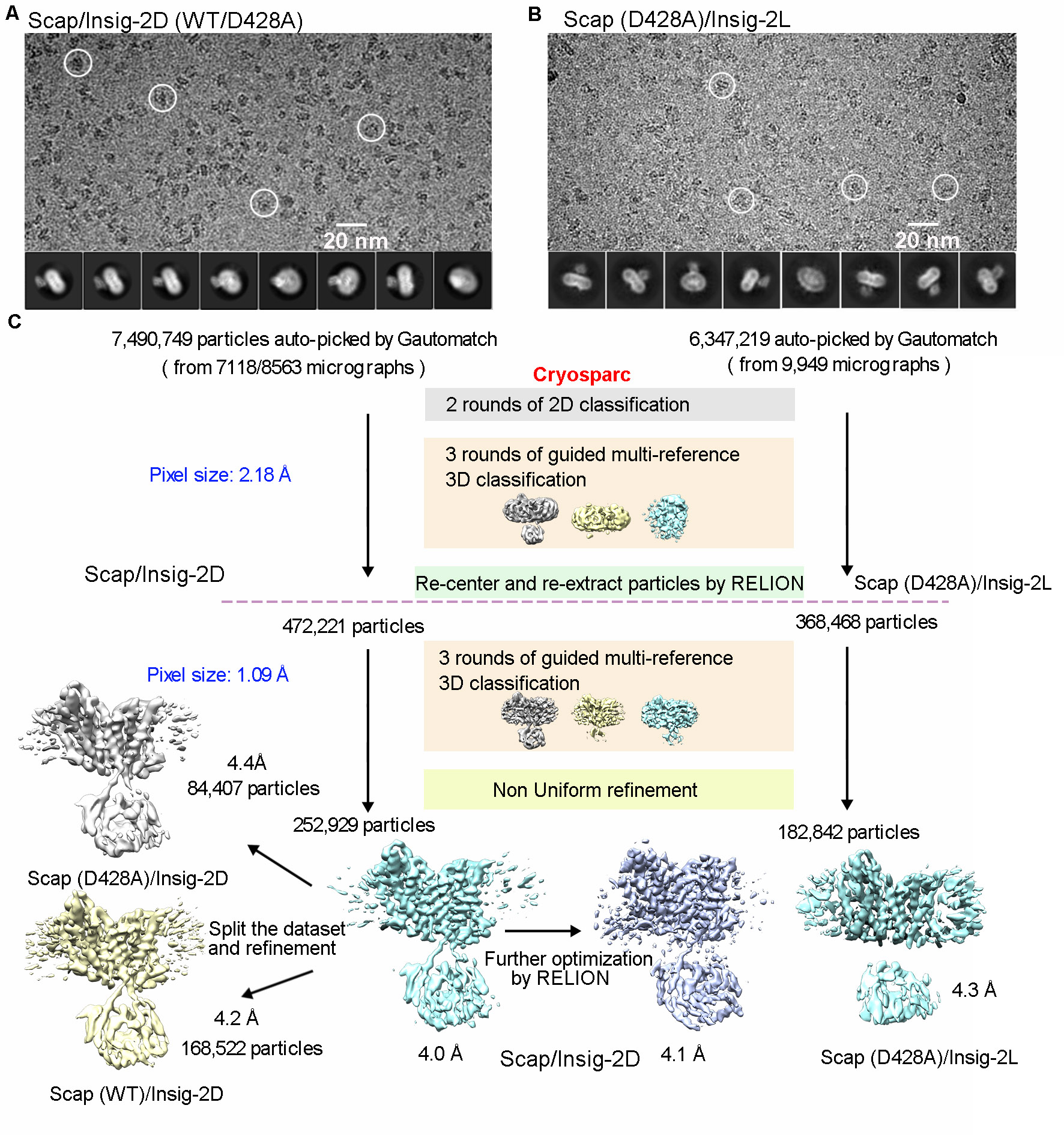

**Fig. S2 | Cryo-EM data processing of the human Scap/Insig-2 complexes.**

(**A**) Representative electron micrograph and 2D class averages for Scap/Insig complexes, both WT and D428A, in digitonin micelles. The data were initially combined for particle picking and separated in the last step as shown in the flowchart in panel C. (**B**) Representative electron micrograph and 2D class averages for Scap (D428A)/Insig complex purified in LMNG. (**C**) The flowchart for EM data processing. Details can be found in Experimental procedures.

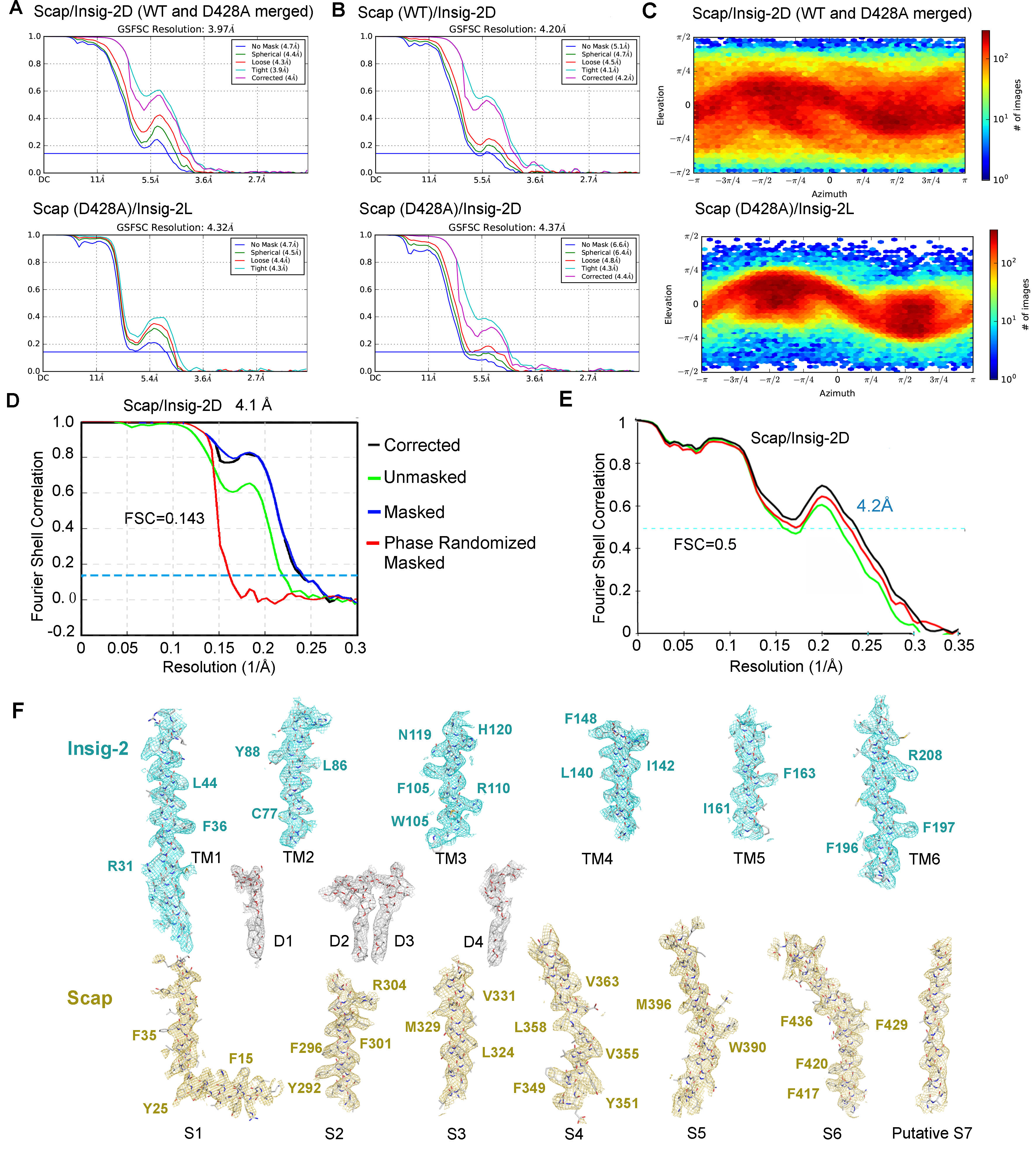

**Fig. S3 | Cryo-EM analysis of the Scap/Insig-2 complexes.**

(**A**) Gold standard Fourier shell correlation (FSC) curves. Shown here are the FSC curves for the 3D refinement of the structures of Scap/Insig-2D (*upper*) and Scap/Insig-2L complexes (*lower*). (**B**) Gold standard Fourier shell correlation (FSC) curves. Shown here are the FSC curves for the 3D refinement of the structures of Scap (WT)/Insig-2D and Scap (D428A)/Insig-2L complexes (*lower*). (**C**) Angular distribution of the particles used for the reconstruction of Scap/Insig-2 complex purified under two conditions. (**D**) Gold standard Fourier shell correlation (FSC) curves of Scap/Insig-2D by RELION. (**E**) FSC curves of the refined TM structure of Scap/Insig-2D versus the TM region map that it is refined against (black); of the model refined against the first half map versus the same map (red); and of the model refined against the first half map versus the second half map (green). The small difference between the red and green curves indicates that the refinement of the atomic coordinates did not suffer from overfitting.

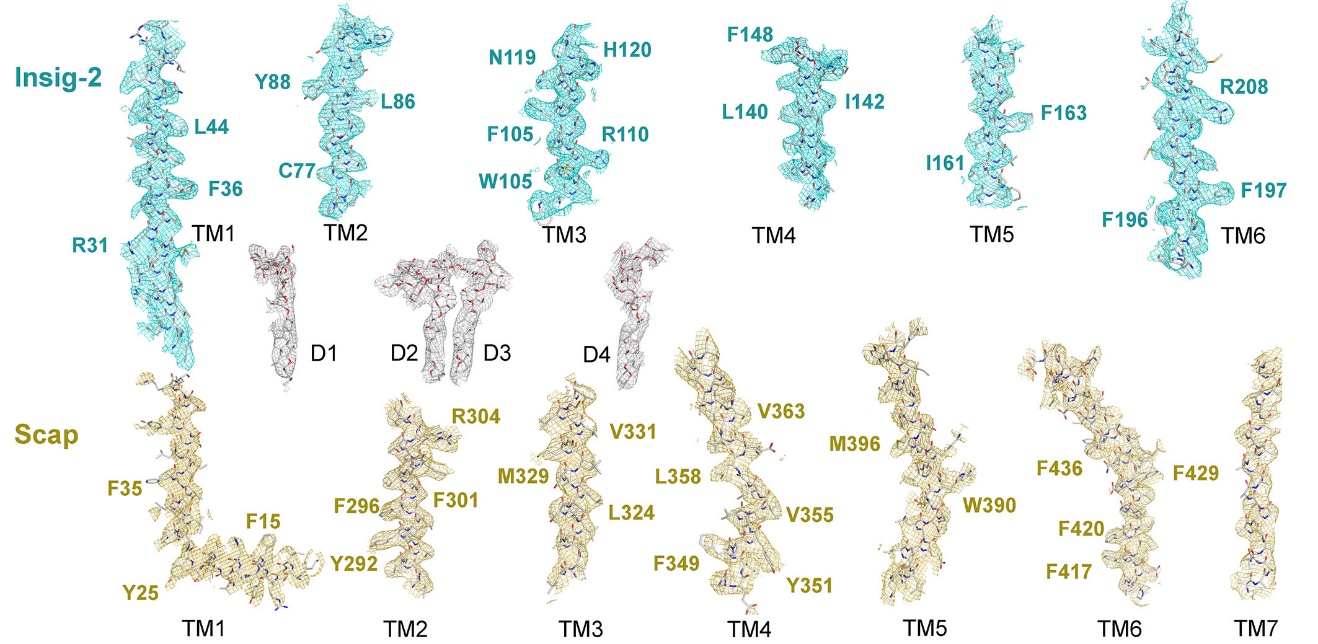

**Fig. S4 | Maps for representative segments in Scap/Insig-2D**.

EM maps for the indicated segments and detergent molecules in Scap/Insig-D. The densities, contoured at 5 σ, were prepared in PyMol.

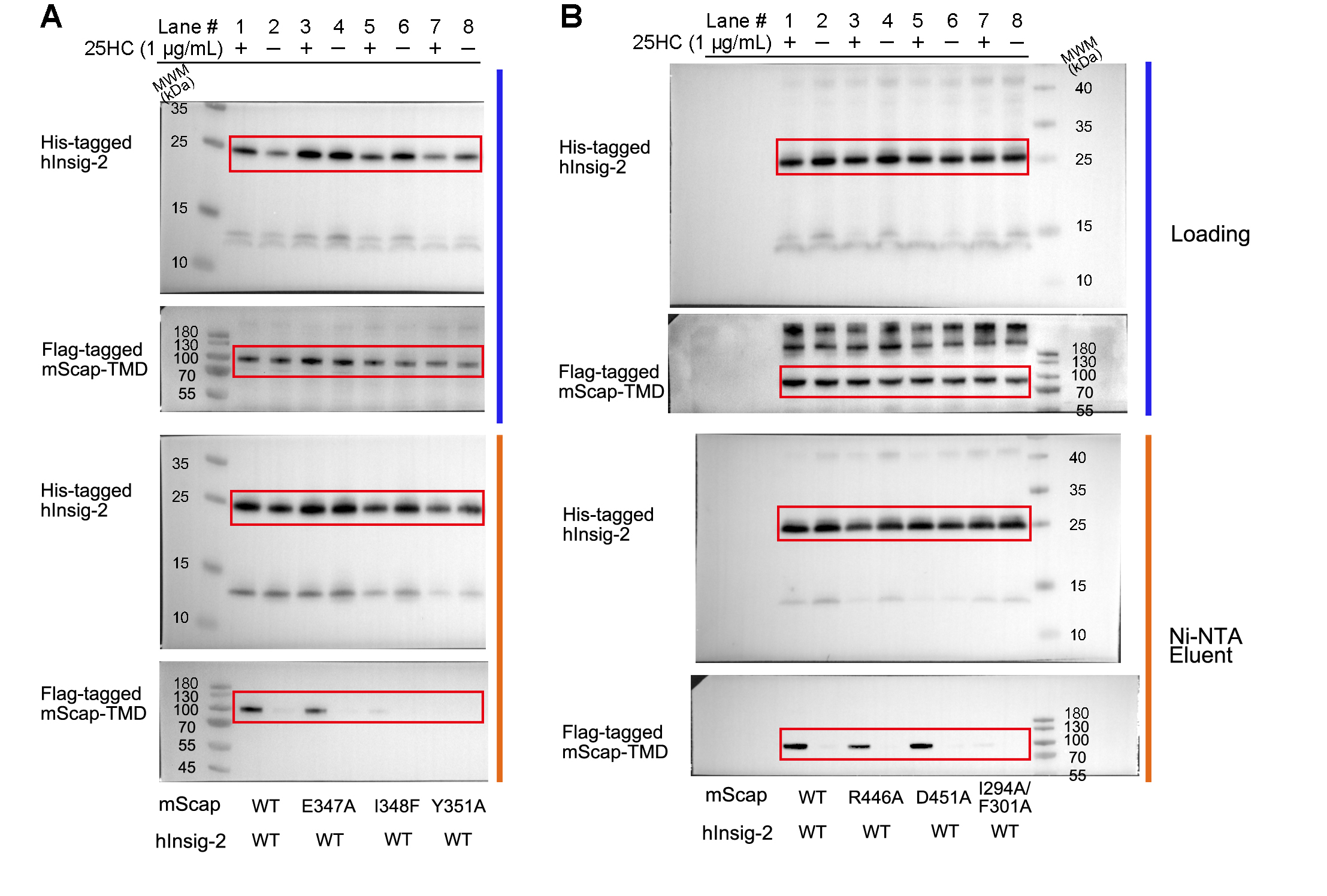

**Fig. S5 | Uncropped blot images of the pull-down assay in Figure 2E.**

The red boxes indicate the presented bands in the main figure. The results shown here and Figure S7 are representative blots from three independently repeated experiments.

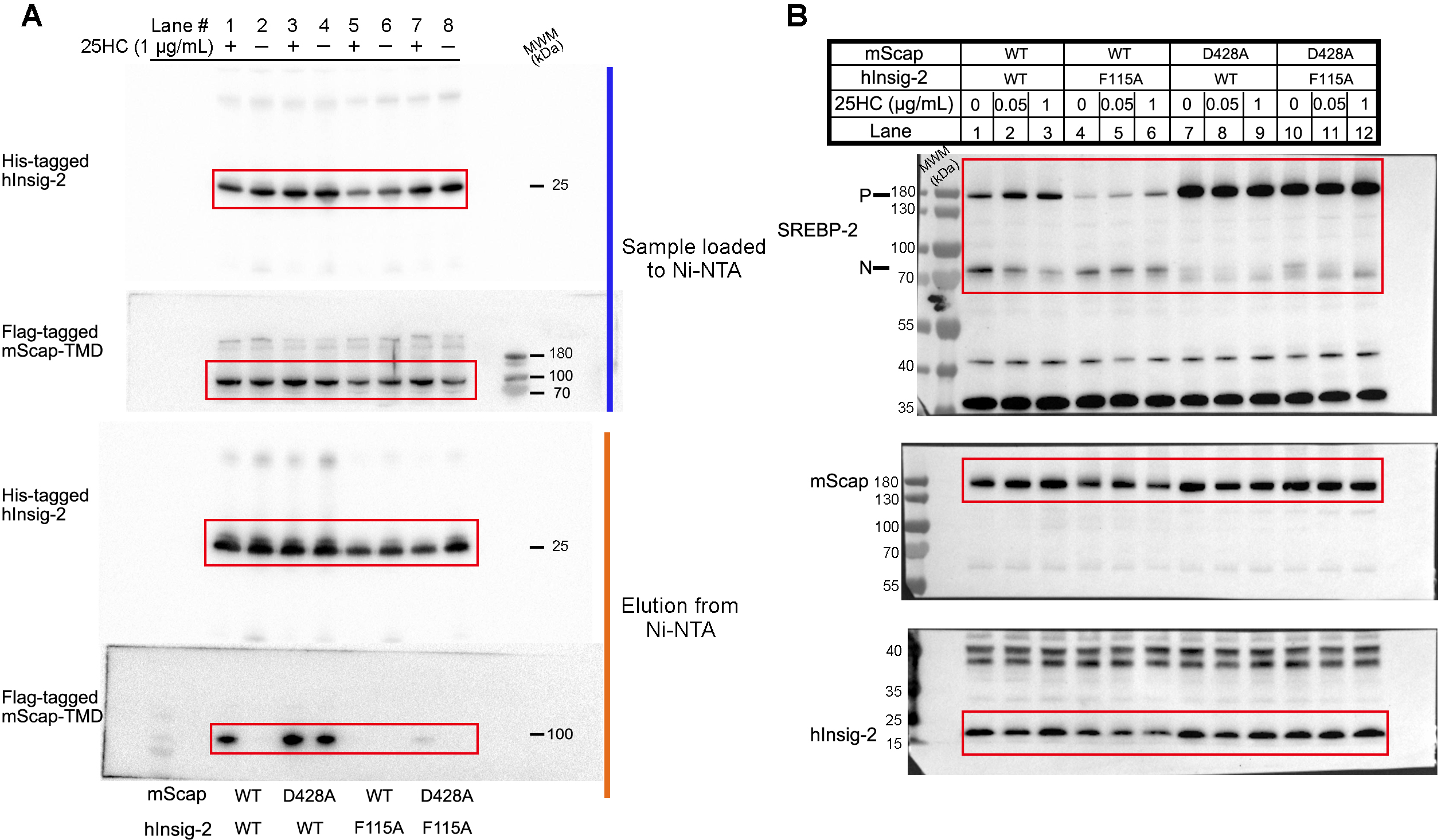

**Fig. S6 | Uncropped blot images of the pull-down assay and SREBP-2 cleavage assays presented in Figure 3.**

(**A**) The blots for Figure 3D. (**B**) Representative blots for Figure 3E. The red boxes indicate the presented bands in the main figures.

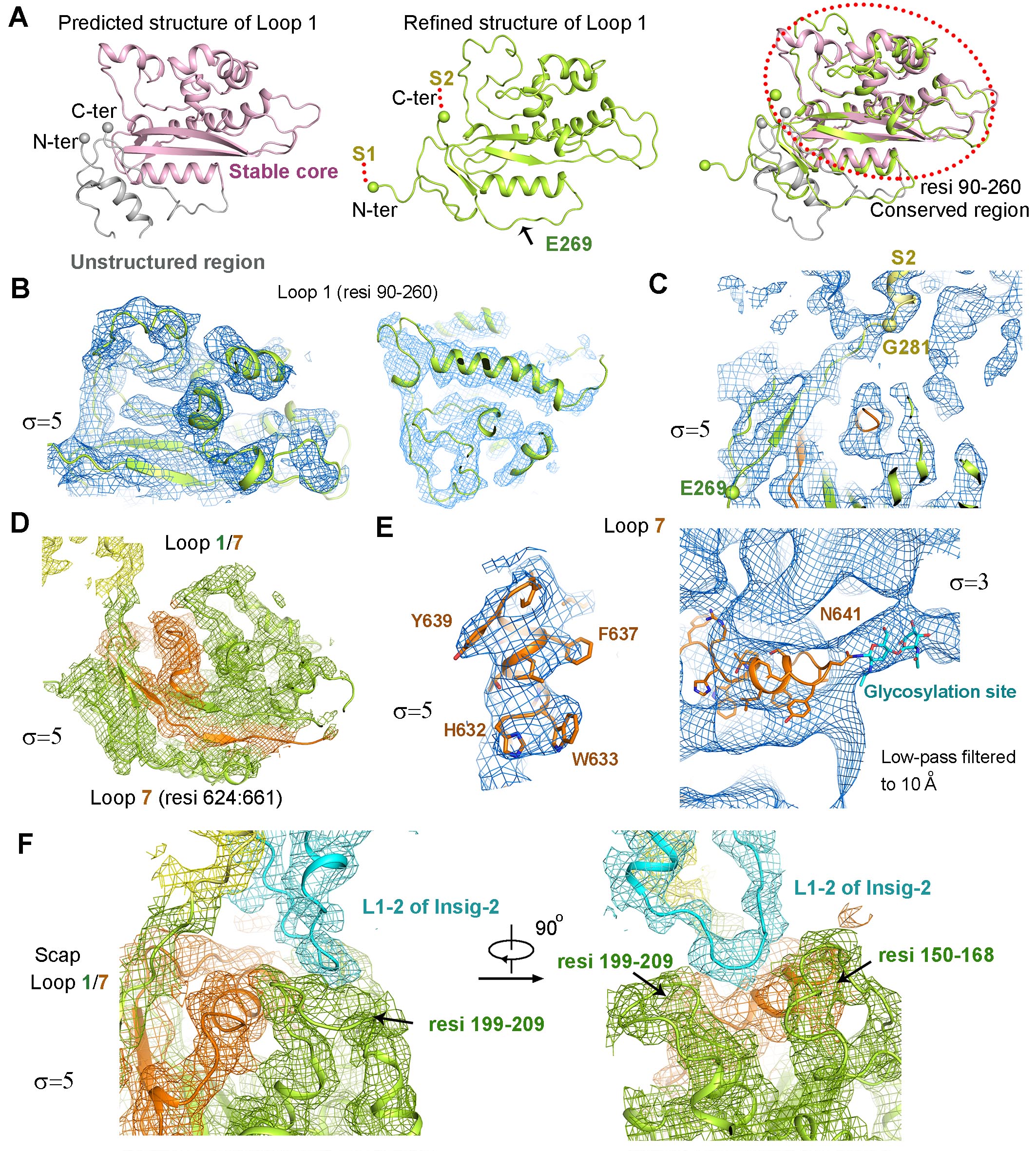

**Fig. S7 |** **AI-facilitated model building for Scap-L1/L7 into the moderate-resolution map.** Please refer to Methods for details.

(**A**) AI-facilitated structural modeling for Scap-L1 I. Shown here are the predicted (*left*), rebuilt and refined (*middle*), and an overlay of the predicted and refined structural models of Scap-L1 (*right*). The structural model for residues 269-281, which exhibit an extended conformation, was grafted directly from the predicted one without further adjustment. The spheres indicate the Cα atoms of the residues that mark the boundaries of the predicted and refined L1 structural model. (**B**) EM map for Scap-L1 contoured at the indicated sigma level. (**C**) The density for the C-terminus of L1 is contiguous with that of the transmembrane S2 segment, validating the sequence assignment. (**D**) EM map for a segment of Scap-L7. (**E**) Sequence assignment of L7 was facilitated by a string of bulky residues (*left*) and glycosylated Asn641 (*right*). (**F**) Simultaneous interactions of the L1-2 loop of Insig-2 with both L1 and L7 of Scap.

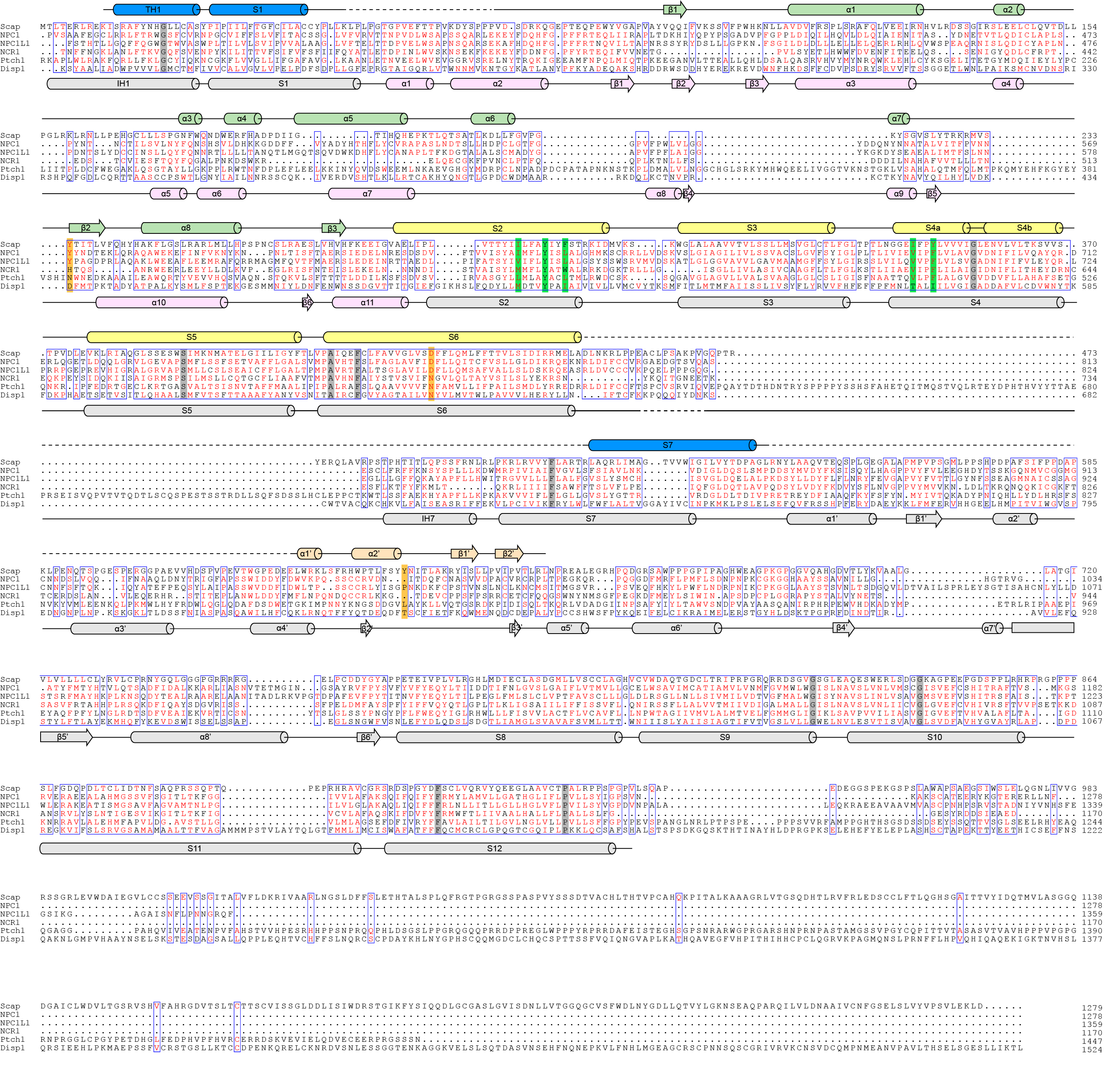

**Fig. S8 | The sequence similarity between Scap-L1/L7 and the luminal and extracellular domains of other SSD-containing proteins was not evident until the structures were revealed.**

Full-length human Scap was aligned with the indicated SSD-containing proteins using the program MultAlin (*16*). Secondary structural elements of Scap and NPC1 are shown above and below the sequences, respectively. The residues with LOF mutations (Y298C, I348F, Y351A, and I294A&F301A) of Scap are shaded green and GOF mutations (Y234A, D428A, and Y640S) are shaded orange. The Uniprot IDs for the proteins aligned are: Scap: Q12770; NPC1: O15118; NPC1L1: Q9UHC9; NCR1 (NPC intracellular sterol transporter 1-related protein 1): Q12200; Ptch1 (Patched 1): Q13635; Disp1 (Dispatched 1): Q96F81.

**Table S1. Cryo-EM data collection and refinement statistics.**

| **Data collection (Scap/Insig-2D)** / **(Scap(D428A)/Insig-2L)** | |
| --- | --- |
| EM equipment | FEI Titan Krios |
| Voltage (kV) | 300 |
| Detector | Gatan K2/K3 |
| Pixel size (Å) | 1.091/1.087 |
| Electron dose (e-/Å^2^) | 45.6/45.6 |
| Defocus range (μm) | 1.5~2.0 |
| **Reconstruction** | |
| Software | RELION 3.0&CRYOSPARC |
| Number of used Particles | 252,929/182,842 |
| Symmetry | C1/C1 |
| Map sharpening B-factor (Å^2^) | -200.0/-200.0 |
| Final Resolution (Å) | 4.1/4.3 |
| **Model building and refinement** | **Scap/Insig-2D** |
| Software | PHENIX & COOT |
| Cell dimensions |  |
| a=b=c (Å) | 218.20 |
| α=β=γ (˚) | 90 |
| Protein residues | 681 |
| Digitonin | 4 |
| R.m.s deviations | |
| Bonds length (Å) | 0.005 |
| Bonds Angle (˚) | 1.156 |
| Ramachandran plot statistics (％) | |
| Preferred | 92.40 |
| Allowed | 6.65 |
| Outlier | 0.95 |
| Molprobity score | 2.53 |
